# Metabolomics and lipidomics in *C. elegans* using a single sample preparation

**DOI:** 10.1101/2020.07.06.190017

**Authors:** Marte Molenaars, Bauke V. Schomakers, Hyung L. Elfrink, Arwen W. Gao, Martin A.T. Vervaart, Mia L. Pras-Raves, Angela C. Luyf, Reuben L. Smith, Mark G. Sterken, Jan E. Kammenga, Antoine H. C. van Kampen, Georges E. Janssens, Frédéric M. Vaz, Michel van Weeghel, Riekelt H. Houtkooper

## Abstract

Comprehensive metabolomic and lipidomic mass spectrometry methods are in increasing demand, for instance in research related to nutrition and aging. The nematode *C. elegans* is a key model organism in these fields, due to the large repository of available *C. elegans* mutants and their convenient natural lifespan. Here, we describe a robust and sensitive analytical method for the semi-quantitative analysis of >100 polar (metabolomics) and >1000 apolar (lipidomics) metabolites in *C. elegans*, using a single sample preparation. Our method is capable of reliably detecting a wide variety of biologically relevant metabolic aberrations in, for instance, glycolysis and the TCA cycle, pyrimidine metabolism and complex lipid biosynthesis. In conclusion, we provide a powerful analytical tool that maximizes metabolic data yield from a single sample.

**Graphical abstract:** 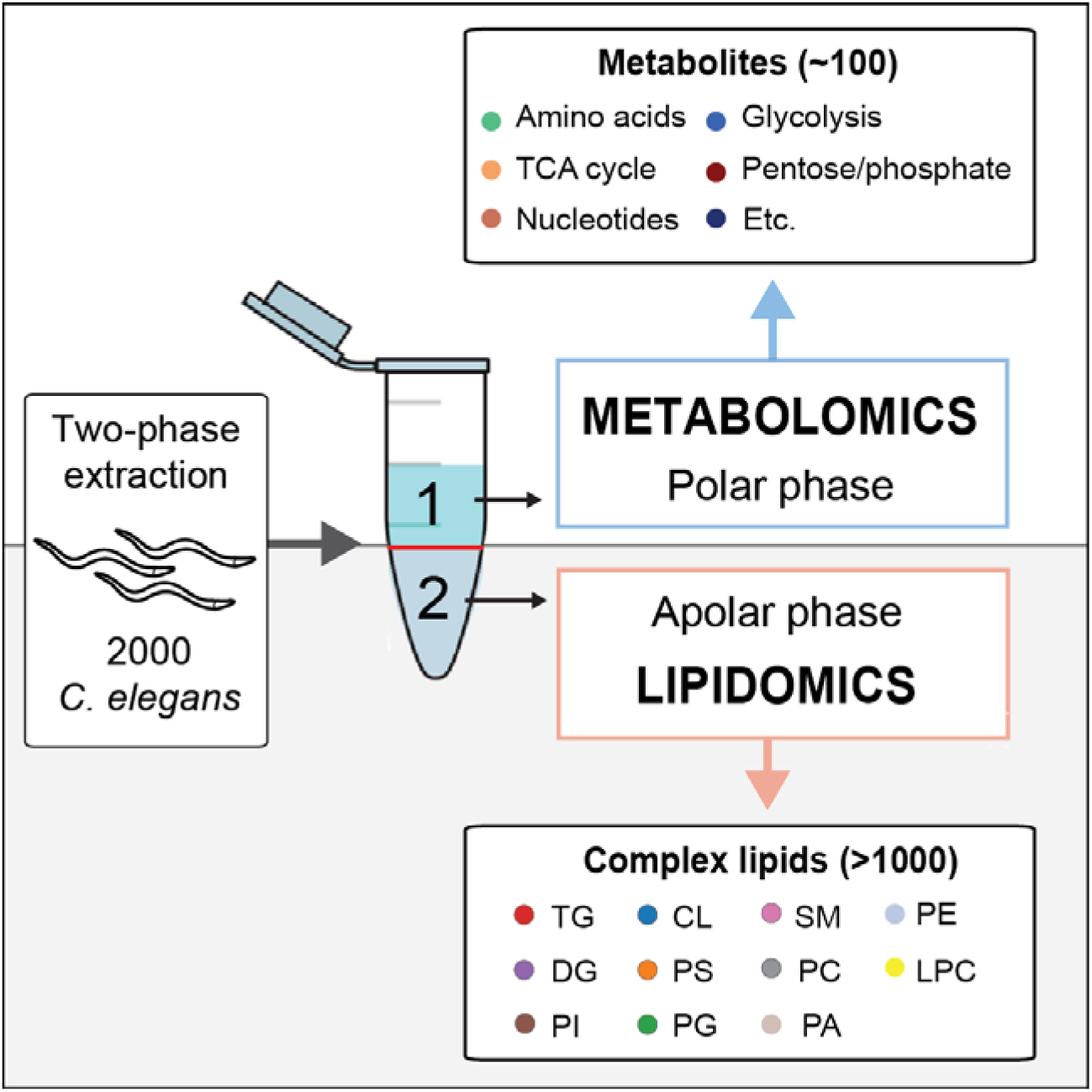

## Introduction

Considerable advances in high-performance liquid chromatography (HPLC), mass spectrometry (MS), nuclear magnetic resonance (NMR) make it possible to reliably detect tens of thousands of compounds^1^. Additionally, semi-automatic annotation of metabolites and data analysis tools have greatly improved the quality and robustness of metabolomic platforms, allowing for an improved sample throughput and ease of data analysis and interpretation^2^. As a consequence, metabolomic analysis has seen a surge in popularity over the last decades and the importance and intricacies of metabolism in health and disease are becoming increasingly evident^2^. In turn, this has prompted increased demand for reliable and robust metabolomic methods for polar and apolar metabolite analyses in model organisms and human tissues^3^.

For many years, *Caenorhabditis elegans* nematodes have been used intensively to investigate genetics, development, as well as aging. *C. elegans* is a versatile model system as genetic influences can be tested with relative ease due to the availability of large repositories of mutants as well as RNAi libraries. Moreover, genetic reference populations have been generated for *C. elegans* in which natural genetic variation is present at a level similar to the human population^4^. This way, meaningful data on population genetics and gene-by-environment interactions can be obtained using for instance dietary interventions^5,6^. More recently, *C. elegans* has become a relevant model to investigate metabolism, since metabolism was identified as a key regulator of traits such as aging^7-9^. Metabolic network models for *C. elegans* were recently constructed^10,11^ and a curated consensus is currently being assembled in a European-led consortium^12^. The success of such endeavors relies heavily on accurate and robust metabolomics methods^13^.

Metabolite measurements in mammalian tissues are commonplace^1,14,15^, however, in *C. elegans* they are sparsely applied^16^. Methods for *C. elegans* metabolite analyses are predominantly based on gas chromatography-MS (GC-MS)^17,18^ and nuclear magnetic resonance (NMR) spectroscopy^16,19^. Drawbacks of these approaches include the need for large quantities of worms and a limited number of metabolites that can be quantified (Table 1). Recent developments using targeted metabolomics with LC-MS allow for the measurement of hundreds of metabolites, including fatty acids and amino acids, in a sample of around 2,000 worms^20,21^.

**Table 1.** Comparison of commonly used metabolomics methods for *C. elegans*.

| **Type of approach** | **Metabolites detected** | **Metabolite type** | **Required # of worms** | **Year** |
| --- | --- | --- | --- | --- |
| GC-MS | ~100 | Polar | 120,000 | 2013<sup>38</sup> |
| GC-MS | 186 | Polar | Unknown | 2015<sup>39</sup> |
| NMR | 45 | Polar | ~8,000 | 2017<sup>40</sup> |
| NMR | 17 | Polar | 200,000 | 2019<sup>41</sup> |
| LC-MS | 18 | Polar<br>(Amino Acids) | 2,000 | 2017<sup>20</sup> |
| LC-MS | 105 | Polar | 2,500 | 2019<sup>21</sup> |
| LC-MS | 82 | Apolar<br>(Spingolipids) | 750 mg<br>(no amount specified) | 2019<sup>42</sup> |
| LC-MS | Untargeted<br>(~3000 features) | Apolar (Lipids) | 8,000 | 2019<sup>43</sup> |
| LC-MS | 44 | Apolar<br>(Fatty Acids) | 2,500 | 2017<sup>20</sup> |
| LC-MS | ~600 | Apolar<br>(Phospholipids) | 2,000 | 2017<sup>20</sup> |
| LC-MS | >1100 | Polar + Apolar | 2,000 | This protocol |

Although these methods are useful when focusing on specific metabolite classes, they rely on separate extraction procedures for both polar and apolar metabolites in biological replicates making it less suitable for screening purposes. Hence, we set out to develop an easy extraction and measurement protocol, allowing for the analysis of a broad panel of polar (metabolome) and apolar (lipidome) metabolites. We shall refer to these as metabolomics and lipidomics respectively. The method presented here provides a detailed step-by-step protocol for sample collection and processing, metabolite extraction, annotation, and relative quantification in *C. elegans.* We demonstrate that metabolomic and lipidomic analysis can be performed on a single sample using a single extraction protocol, reducing sample preparation and throughput time without compromising metabolite identification. With this protocol we can semi-quantitatively measure >100 polar and >1000 apolar metabolites, from all major metabolite classes in a sample of approximately 2000 worms. Moreover, this method can be easily adapted for other model systems, cells, and tissues.

## Results

### Combined extraction for polar and apolar metabolites

Endogenous metabolites span a wide range of physicochemical properties, making it difficult to extract a large range of the metabolome with a single solvent. Additionally, polar solvents typically lack the ability to precipitate interfering proteins in biological samples, making a simple water extraction of polar metabolites impractical. An elegant way to remedy this issue, is to use a liquid-liquid extraction. In this case, a highly apolar solvent, e.g. chloroform, is used to precipitate protein and facilitate the breakdown of biological organization, while a polar solvent is added to extract polar metabolites in a separate layer. This type of extraction also doubles as a separation step, removing apolar compounds from the polar layer (and vice versa), thereby reducing ion suppression effects during MS analysis. Interestingly, this type of two-phase extraction was first applied for extracting lipids in the (predominantly) chloroform phase^22^. We used such a two-phase extraction on *C. elegans* to perform both metabolomics and lipidomics in a single sample. Combining the analysis of metabolomics and lipidomics allows for the analysis of a broad spectrum of metabolites in a single extraction step, requiring only a single sample. This way, the required number of *C. elegans* cultures is halved, waste is significantly reduced, and comparisons between the two omics analyses are more meaningful.

### Validation of polar metabolite (metabolomics) analysis in *C. elegans*

In order to enable validation, we used *C. elegans* pellets containing different numbers of worms and extracted polar metabolites from the upper phase of the liquid-liquid extraction (Fig. 1a). Polar metabolites were separated using a variation of Hydrophilic Interaction Liquid Chromatography (ZIC-cHILIC) and measured using a Q Exactive Plus Orbitrap mass spectrometer. For each of the annotated polar metabolites, we determined their linear response since loss of linearity of the MS response is a good measure of bias in sample preparation or analysis (Supplementary Table 1). Four example metabolites, i.e. pyruvate, cytidine monophosphate (CMP), adenosine triphosphate (ATP), and nicotinamide adenine dinucleotide (NAD^+^) show strong linearity across the range of worms before applying normalization for internal standard (Fig. 1b-e, Supplementary Table 1). We then established which internal standards to use for each metabolite (Supplementary Table 2). Selection of the internal standard was based on the combination of the lowest coefficient of variation for duplicates, and highest Pearson correlation coefficient after correction across the number of worms (Fig. 1f-i). Applying these internal standards for the data normalization led to even better linearity of for pyruvate, CMP, ATP, and NAD^+^ (Fig. 1j-m, Supplementary Table 2). Altogether, we established optimal internal standard conditions for each of the polar metabolites and found that each of these can be reliably measured in a sample of ∼2000 worms.

**Fig. 1.**
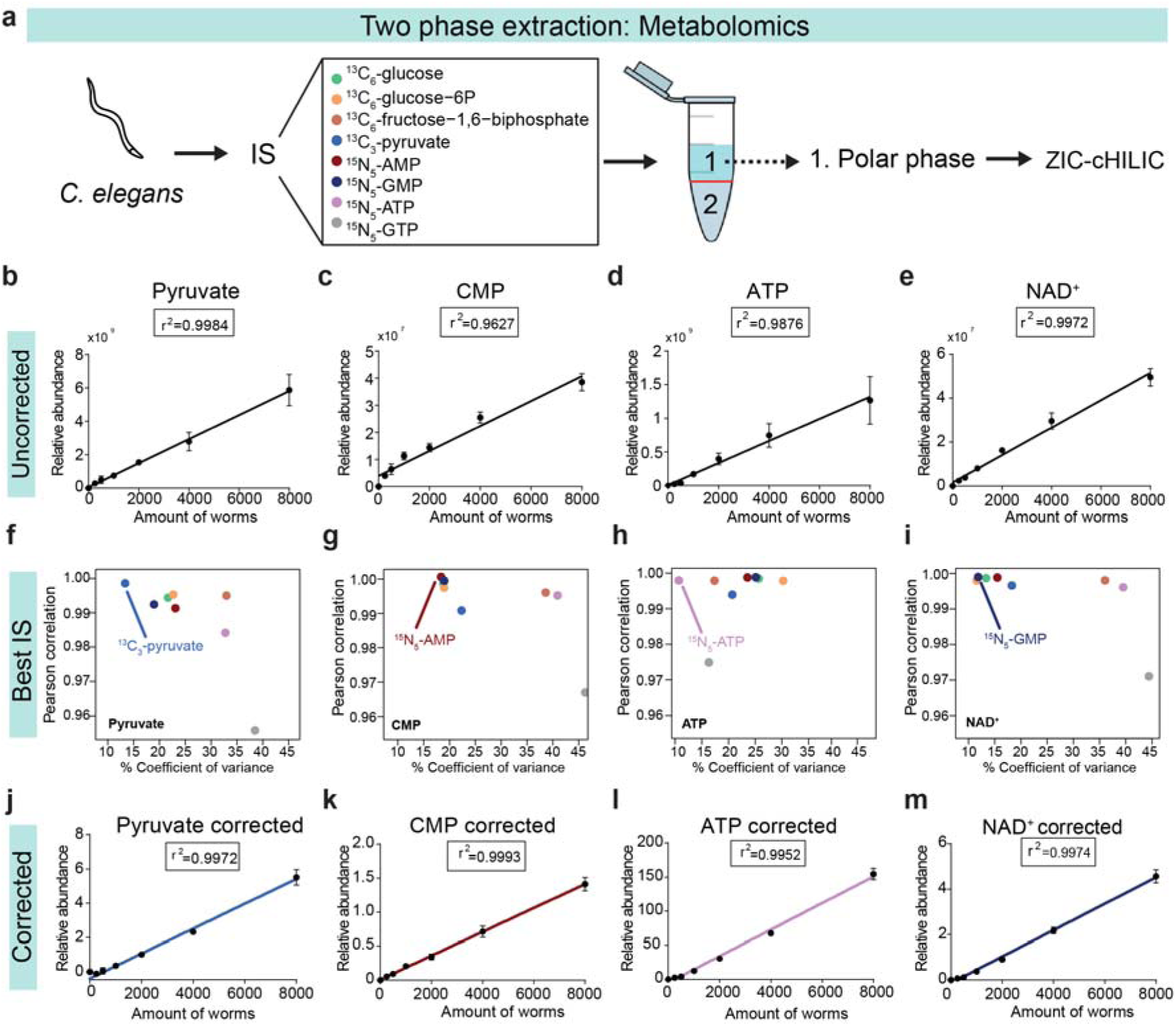
Validation and linearity of metabolites extracted from polar phase. **a**, Internal standard (IS) was added to *C. elegans* pellet and using a two-phase extraction the upper polar phase was processed for ZIC-cHILIC. Linearity of four example metabolites **b**, Pyruvate, **c**, CMP, **d**, ATP, and **e**, NAD^+^ shows r^2^ >0.98. For **f**, Pyruvate, **g**, CMP, **h**, ATP, and **i**, NAD^+^ the best IS was determined per metabolite by plotting Pearson’s correlation coefficient against coefficient of variance. Linearity of **j**, Pyruvate **k**, CMP **l**, ATP, and **m**, NAD^+^) after correction for their best IS shows r^2^ >0.99. Data points represent mean +/- SD with n=4.

### Validation of the apolar metabolite (lipidomics) analysis

Instead of a dedicated single-phase extraction we reported before^23^, we now used the “left-over” apolar phase from our two-phase extraction to analyze lipids (Fig. 2a, Supplementary Table 3). Lipids were separated using both a normal-phase (NP) and reversed-phase (RP) chromatography method and measured on a Q Exactive Plus mass spectrometer. The lipidome is an enormously diverse class of metabolites with widely varying polarities. For instance, at the apolar end of the spectrum, triacylglycerols (TGs) consist of a glycerol backbone and three fatty acids tails. Due to these uniformly apolar qualities, TGs of any chain length partition almost exclusively to the chloroform phase during the two-phase extraction. This results in a linear relationship between the number of worms and the measured abundance of TGs (Fig 2b). A similar pattern is observed for other major lipid classes containing multiple acyl side chains, such as diacylglycerols (DGs), phosphatidylinositols (PIs), cardiolipins (CLs), phosphatidylserines (PSs), and phosphatidylglycerols (PGs) (Fig. 2c-g). Sphingomyelins (SMs) have a different basic structure, but also contain two alkyl moieties (the sphingosine backbone and the N-acyl group), resulting in good linearity (Fig. 2h). Finally, while the most abundant PL class phosphatidylcholines (PCs) has two acyl groups and is extracted in the apolar phase, it has a lower r^2^ of 0.667 (Fig. 2i). This is due to its high abundance, saturating the detector at higher worm numbers. Indeed, when considering linearity for ≤4000 worms, the r^2^ for PC is 0.832 and even goes up to r^2^=0.902 when analyzing up to 2000 worms. It is therefore advised to use ≤4000 worms in order to accurately measure PCs. Phosphatidylethanolamines (PEs) show the same trend though to a lesser extent (Fig. 2j).

**Fig. 2.**
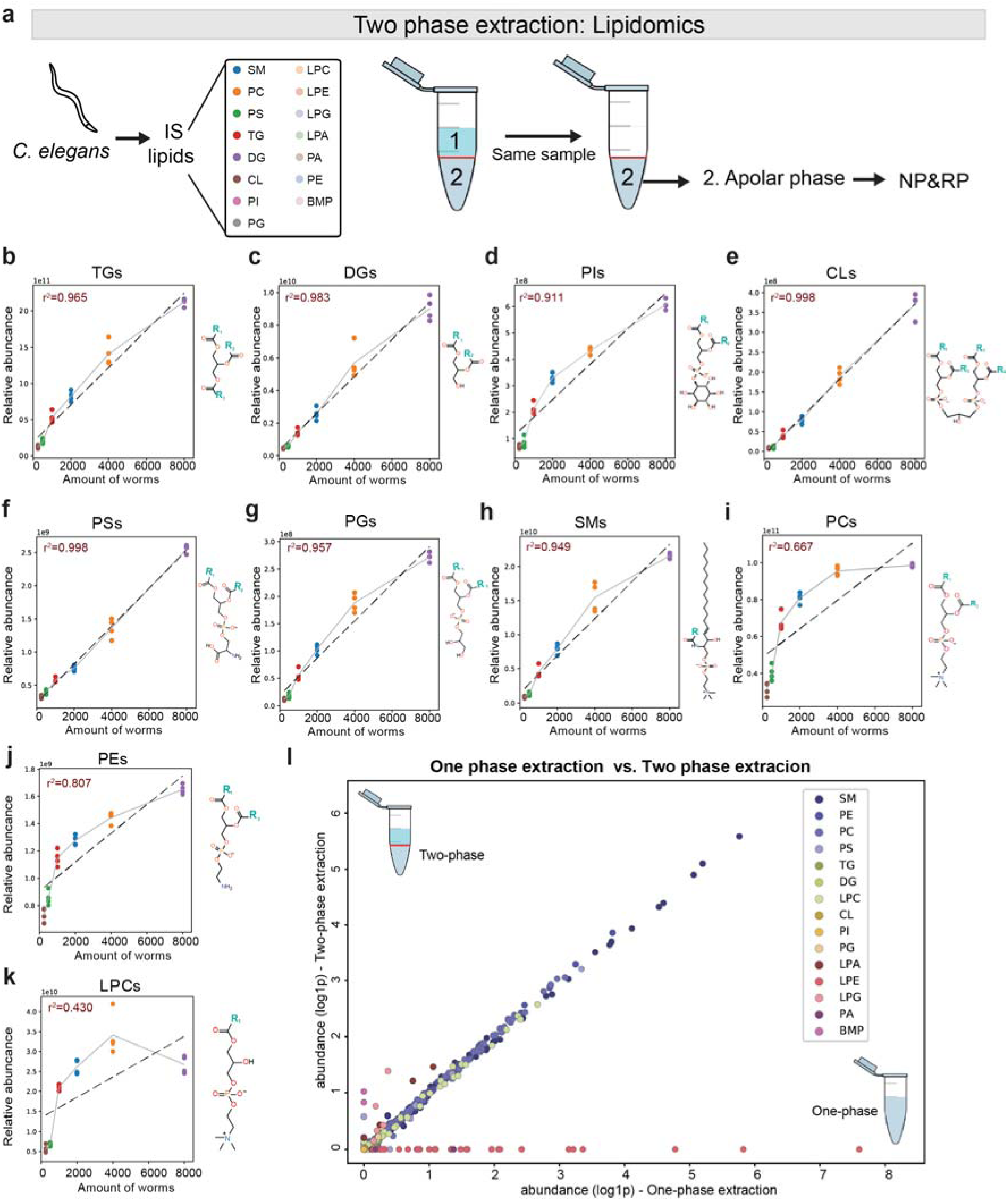
Validation of lipids extracted from apolar phase and comparison of two-phase vs. one-phase lipid extraction. **a**, IS was added to *C. elegans* pellet and using a two-phase extraction the lower apolar phase from the same sample was processed for lipidomics normal phase (NP) and reversed phase (RP). Linearity of **b**, TGs **c**, DGs **d**, PIs **e**, CLs **f**, PSs **g**, PGs **h**, SMs **I**, PCs **j**, PEs and **k**, LPCs **l**, Comparison of relative abundances (log1p) from one-phase extraction vs. two-phase extraction.

At the other end of the lipid polarity spectrum are lysolipid species such as lysophosphatidylcholine (LPC), lysophosphatidic acid (LPA), lysophosphatidylethanolamine (LPE), and lysophosphatidylglycerol (LPG), each containing only a single fatty acid side chain (R_1_), and a polar head group. Similar to PC, LPC abundance is high and its detection reaches a plateau at higher worm numbers (Fig. 2k). When including all data points, up to 8000 worms, the r^2^ is 0.430, but this improves (r^2^ = 0.872) when including ≤2000 worms. Other lyso-phospholipids are poorly detected in chloroform phase of the current two-phase method, resulting in loss of linearity (Fig. S1a-c). Interestingly, despite containing two fatty acid side chains, phosphatidic acid (PA), and bis(monoacylglycero)phosphate (BMP) show r^2^’s of 0.785 and 0.667, respectively (Supplemental Fig. 1d-e). Due to these complex lipid properties relating to solubility, it is likely that these lipids are (partly) extracted to the polar phase during the two-phase liquid-liquid extraction.

To determine the extent of these partitioning effects on the detected lipidome, we made a direct comparison between the detected lipidome of the one-phase extraction^20^ and the new two-phase extraction of samples containing ∼2000 worms from the exact same biological experiment. When plotting all the individual lipid species, we observed that for most lipids the measured abundance is highly similar between the one-phase and the two-phase extraction (Fig. 2l), suggesting that there was no significant loss of these lipid species in the polar extraction phase. However, around 9% of the species that are normally detected using the one-phase extraction are not detected when applying the two-phase method, most strikingly the entire LPE class (Supplemental Fig.1b). On the other hand, 10% of the lipids were only detected in the two-phase extraction, such as some BMPs and other low-abundant lipids (Fig. 2l). Possibly, these low-abundant lipids are not detected with the one-phase extraction due to suppression effects in the MS. In conclusion, despite the loss of some polar lipid species, including the whole LPE class, there were some low abundant lipid species only recovered using the two-phase extraction. Most importantly, the vast majority of major lipid classes are detected equally well with the two-phase extraction compared to the one-phase extraction.

### Polar and apolar metabolites change upon knockdown of metabolic genes in *C. elegans*

In order to test how well our method was able to pick up biologically relevant differences, we first targeted four different metabolic pathways using RNAi against either enzymes or other factors known to affect metabolic pathways in *C. elegans* (Supplementary Table 4).

The first gene targeted was the pyruvate dehydrogenase alpha subunit (*pdha-1*), which is part of the complex that is responsible for converting pyruvate into acetyl-CoA. Since acetyl-CoA feeds into the TCA cycle it links glycolysis to the TCA cycle, and RNAi of this enzyme is expected to affect both of these pathways. Indeed, worms treated with *pdha-1* RNAi show many significant changes in metabolite abundance (Fig. 3a). A five-fold increase was observed for pyruvate in these worms compared to worms treated with an empty vector, and a two-fold increase was observed for alanine, which is an amino acid that can be formed from pyruvate (Fig. 3b). On the other hand, a significant decrease was observed for all TCA cycle intermediates we measured, including: (iso)citrate, α-ketoglutarate, succinate, fumarate, malate and oxaloacetate (Fig. 3b). This is in line with a reduced availability of acetyl-CoA that can enter the TCA cycle.

**Fig. 3.**
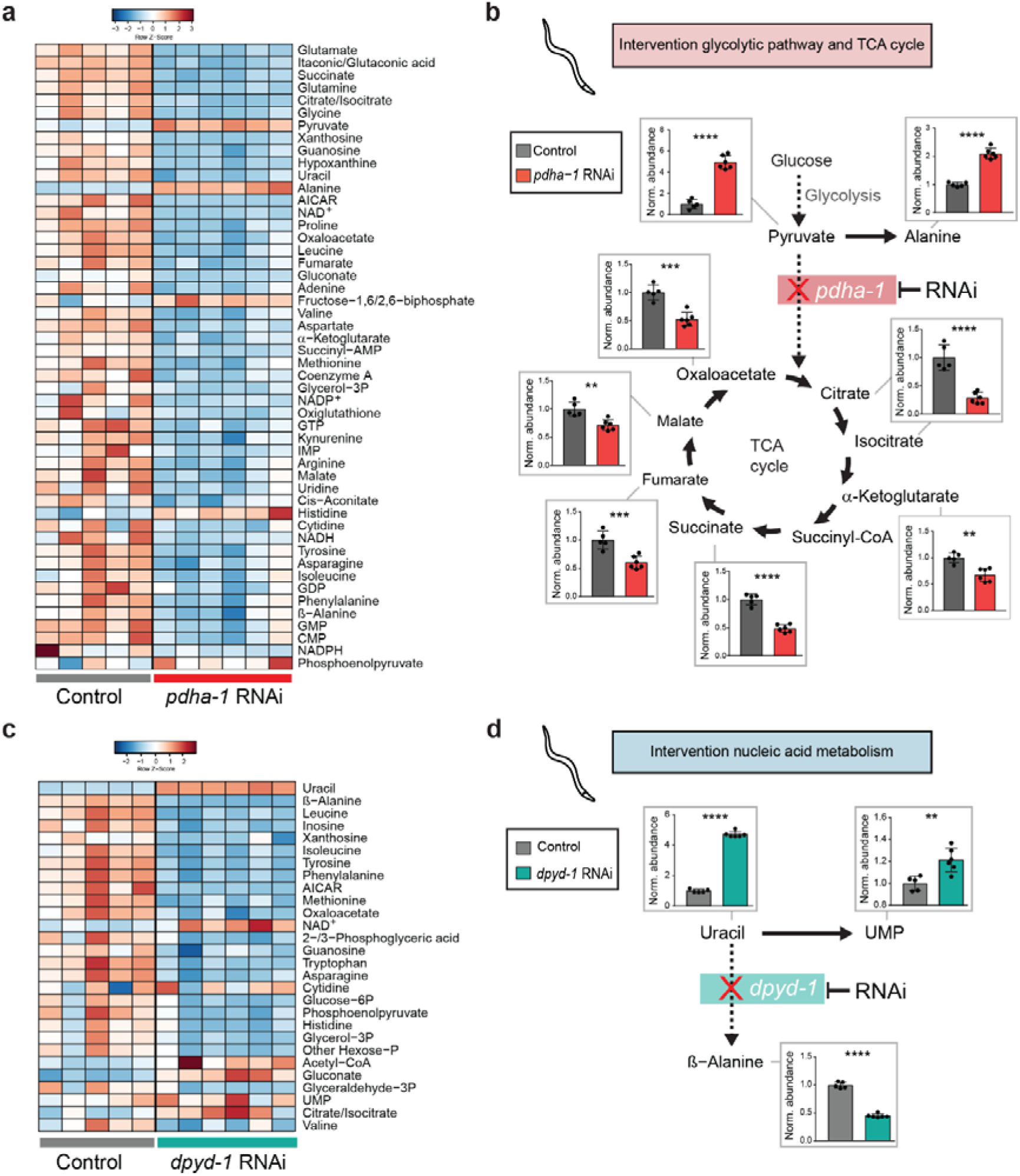
Metabolite changes in worms treated with either *pdha-1* RNAi or *dpyd-1* RNAi. **a**, Heatmap of metabolite changes sorted on FDR of *pdha-1* RNAi treated worms compared to control worms treated with an empty vector. **b**, RNAi of the *pdha-1* enzyme metabolizing pyruvate into acetyl-CoA, providing the primary link between glycolysis and the tricarboxylic acid (TCA) cycle, results in significant increases of pyruvate and alanine and significant decrease of TCA cycle intermediates (iso)citrate, α-ketoglutarate, succinate, fumarate, malate and oxaloacetate. **c**, Heatmap of metabolite changes sorted on VIP-score of *dpyd-1* RNAi treated worms compared to control worms treated with an empty vector. **d**, RNAi of the *dpyd-1* enzyme, catalyzing uracil which ultimately ends in β-alanine (via 5,6-dihydruracil and N-carbamyl-β-Alanine), results in significant upregulation of uracil and UMP and significant downregulation of β-alanine.

The next enzyme we targeted was dihydropyrimidine dehydrogenase (*dpyd-1*) involved in pyrimidine base degeneration. DPYD-1 is important for nucleic acid metabolism as it catalyzes the reduction of uracil and is involved in the degradation of the chemotherapeutic drug 5-fluoroacil (5-FU). In this *dpyd-1* RNAi condition we also found many significant metabolite changes compared to worms treated with empty vector (Fig. 3c). We observed an almost five-fold accumulation of uracil, accompanied by a small increase of UMP which can be alternatively metabolized from uracil (Fig. 3c-d). DPYD-1 is also involved in ß-alanine biosynthesis^24^. In line with this, a strong reduction of ß-alanine was observed in the *dpyd-1* RNAi treated worms (Fig. 3c-d) again reflecting the knockdown of this enzyme on the metabolite level.

We next set out to establish biological validation of our lipidomics analysis. With RNAi, we targeted an enzyme involved in fatty acid elongation, *elo-2* (Fig. 4a). When exploring the lipid profile of worms using our method, we could distinguish the control worms from the *elo-2* RNAi-treated worms, as shown with PCA analysis (Fig. 4b). In order to visualize effects on the lipid elongation, we then plotted carbon chain length versus the total number of double bonds in those chains for individual lipid classes. For instance, TGs in *elo-2* RNAi-treated worms showed a marked decrease of lipids with long carbon chains and accumulation of lipids with shorter carbon chains regardless of the number of double bonds (Fig. 4c). The same was observed for PCs, DGs (Fig. 4d-e) and other lipid classes (Supplemental Fig. 2). Our data confirm that elongation of carbon chains in fatty acids is inhibited when *elo-2* is knocked down in *C. elegans*, which leads to widespread changes across the lipidome.

**Fig. 4.**
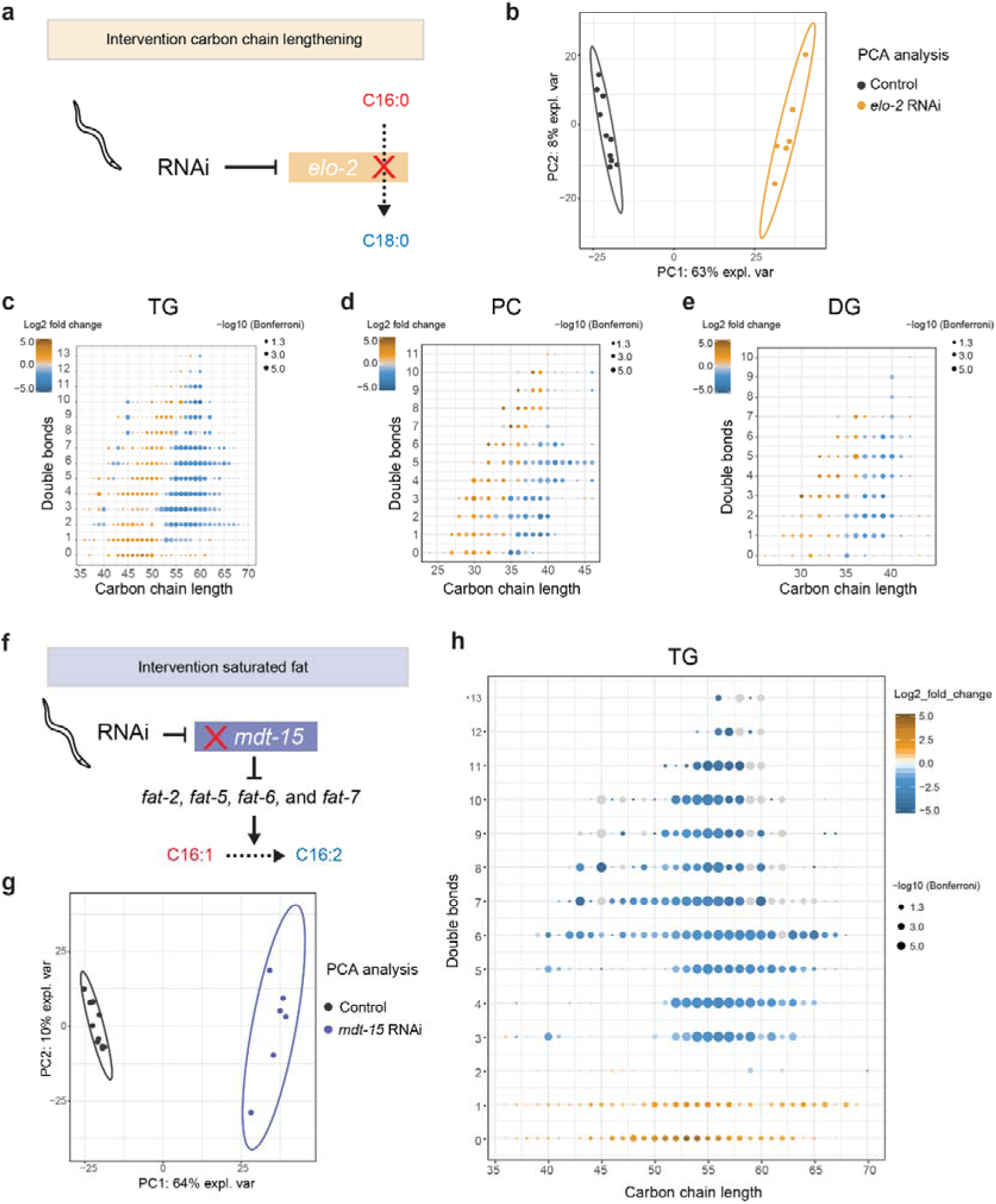
Phospholipid changes in worms treated with either *elo-2* RNAi or *mdt-15* RNAi. **a**, RNAi of *elo-2* which has fatty-acid elongase activity in worms, is expected to increase carbon-chain length of phospholipids. **b**, PCA analysis showing clear distinction between *elo-2* RNAi treated worms and control worms. **c**, Changes in the triacylglycerol (TG) composition of elo-2 RNAi versus empty vector controls shows significant decrease of phospholipids with long carbon chain length (>55) and significant increase of TG species with short carbon chain length (<55). A similar pattern was observed in other phospholipid species such as **d**, PC and **e**, DG. **f**, RNAi of *mdt-15*, a transcription factor upregulating *fat-2, fat-5, fat-6* and *fat-7*, affects the level of unsaturation, i.e. carbon-chain double bonds. **g**, PCA analysis showing clear distinction between *mdt-15* RNAi treated worms and control worms. **h**, Changes in phospholipids of the triacylglycerol (TG) species shows significant decrease of lipids with >2 double bonds and increase of PL with <2 double bonds.

Finally, we targeted *mdt-15*, a subunit of the Mediator complex. Rather than acting on fatty acid elongation, *mdt-15* transcriptionally regulates fatty acid desaturases including *fat-2, fat-5, fat-6* and *fat-7* (Fig. 4f)^25^. The PCA analysis shows clear separation of the control worms from the *mdt-15* RNAi treated worms based on their lipid profiles (Fig. 4g). When plotting carbon chain length versus the number of double bonds, we observed a strong decrease of lipids with multiple double bonds and accumulation of lipids with ≤1 double bond, irrespective of the carbon-chain lengths (Fig. 4h, Supplemental Fig. 3). This shift towards saturated TG species in worms treated with *mdt-15* RNAi is in line with the previously described regulation of *mdt-15* on fatty acid desaturation enzymes^25^.

Together, these four different RNAi conditions affecting distinct metabolic pathways and metabolite classes illustrate that our method can adequately pick up relevant biological differences.

### Metabolic diversity in a *C. elegans* reference population

Our method also allows for the exploration of the natural variation of metabolite abundances occurring due to the differences among genetic backgrounds. To demonstrate this, we turned to recombinant inbred lines (RILs) derived from wild-type worm strains N2 and CB4856^4,26^. RILs are genetic mosaics of the parental strains N2 and CB4856. We reasoned that the sensitivity of our approach could reveal the genome’s more subtle influences due to naturally occurring polymorphisms affecting the metabolome. Therefore, we proceeded to perform the current metabolomics method on two parental wild type strains (N2 and CB4856) and eight different RIL strains resulting from the genetic cross (Fig. 5a, Supplementary Table 5). Metabolic profiling revealed the underlying diversity of metabolites present in the different genetic backgrounds (Fig. 5b). To explore this in a more systematic manner, we calculated the broad-sense heritability (*H*^*2*^) for each metabolite, for both parental and offspring strains^8^. Broad-sense heritability serves as an indication for the percentage of variance for a given metabolite that is explained by genetics. Plotting broad-sense heritability for the parental versus offspring strains illustrates where new combinations of alleles may have severe effects on distinct metabolic profiles and thus indicate genetic complexity of the trait (Figure 4c).

**Fig. 5.**
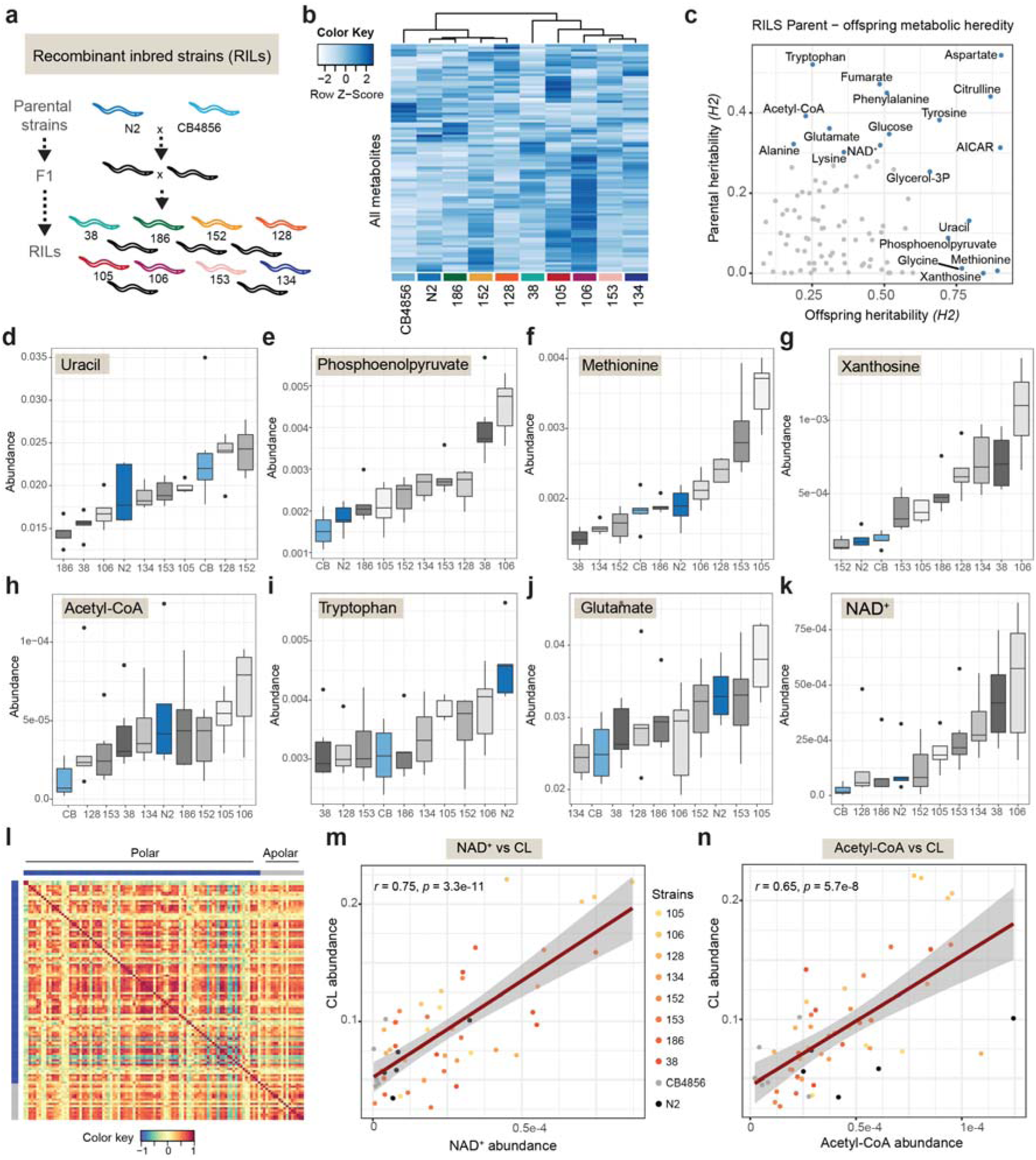
Natural diversity of metabolite abundances in Recombinant Inbred Lines (RILs) **a**, RILs strains selected for analysis **b**, Heatmap showing RILs and metabolites showing diversity in metabolite levels present between the different strains (n=5-6). **c**, broad-sense heritability (*H*^2^) of offspring versus parental lines. Heritability indicates the percentage of variance for a given metabolite that is explained by genetics. Examples diverse heritable outcomes include: **d**, uracil (offspring H2 0.794, FDR < 0.05; parental H2 0.131, FDR not significant) **e**, phosphoenolpyruvate (offspring H2 0.721, FDR < 0.05; parental H2 0.088, FDR not significant) **f**, methionine (offspring H2 0.892, FDR < 0.05; parental H2 0.006, FDR not significant) **g**, xanthosine (offspring H2 0.843, FDR < 0.05; parental H2 0.000, FDR not significant) **h**, acetyl-CoA (offspring H2 0.228, FDR not significant; parental H2 0.392, FDR < 0.05)**i**, tryptophan (offspring H2 0.252, FDR not significant; parental H2 0.521, FDR < 0.05) **j**, glutamate (offspring H20.310, FDR not significant; parental H20.361, FDR < 0.05) **k**, NAD^+^(offspring H2 0.486, FDR not significant; parental H2 0.319, FDR < 0.05). **l**, Cross-correlation matrix between polar (blue) metabolites and apolar (grey) lipid classes, highlighting the range of strong positive correlations (red) to strong negative correlations (blue) between all metabolites and lipid classes. **m**, Example of correlation between apolar cardiolipins (CL) and polar NAD^+^ (Pearson’s r=0.75, p=3.3e-11), parental lines and strains color coded as highlighted in the legend. **n**, example of apolar cardiolipins (CL) and polar acetyl-CoA (Pearson’s r=0.65, p=5.7e-8).

Assessing heritability in this manner, we observed metabolites for which parental strains possessed a low heritability score and their offspring possessed a high heritability score, indicating that there may be multiple loci of opposing effects regulating the metabolite’s abundance. For example uracil, phosphoenolpyruvate, methionine, and xanthosine, exhibited such patterns (Figure 4d-g). Conversely, we also observed metabolites for which the parental strains possessed a high heritability score and the offspring possessed an equal or lower score, which likely indicates that there may be few, or even just a single locus affecting the metabolite’s abundance. Examples of these included acetyl-CoA, tryptophan, glutamate, and NAD^+^ (Figure 4h-k).

Performing metabolomics and lipidomics on the same samples enables data integration from both techniques. To illustrate, we performed cross-correlations between polar metabolites and lipid classes and visualized these in a correlation matrix (Fig. 5l). For example, we identified metabolites that correlated with cardiolipin (CL), which is an important component of the inner mitochondrial membrane^27^. We found that abundance of CL correlated significantly with NAD^+^ (Fig. 5m), which governs mitochondrial function through its role as an enzyme cofactor as well as being a substrate for sirtuins^28,29^. Likewise, CL significantly correlated with acetyl-CoA (Fig. 5n), which is in line with its role in acetyl-CoA synthesis^30^. These correlations are an example of how metabolomics and lipidomics data can be integrated and explored to gain deeper insight into their cross-talk and interrelations.

## Discussion

Changes in metabolism are increasingly recognized as valuable markers of, as well as causal contributors to, the development of metabolic disease and aging. Increasingly comprehensive methods for the analysis of both polar (metabolomics) and apolar (lipidomics) metabolites have proven essential in these fields. Thus far however, these omics methods required dedicated sample preparation, and thus a separate sample, for metabolomics and lipidomics respectively. Here, we report a method that uses both the polar (metabolomics) and apolar (lipidomics) layer of a two-phase liquid-liquid extraction and analyzes these using high resolution MS methods, providing an elegant way of exploring a large range of the metabolome and lipidome in a single sample, covering >1100 annotated metabolites of different classes. We show here that this method is robust and sensitive enough to analyze a wide variety of metabolic pathways using both metabolomics and lipidomics, and capable of reliably pinpointing metabolic aberrations in these pathways.

Biologically relevant differences in central carbon as well as lipid pathways could be determined in detail. The effects of *pdha-1* inhibition using RNAi was reflected throughout the TCA cycle, and the inhibition caused a decrease of acetyl-CoA and an accumulation of its precursor, pyruvate. Additionally, the pyruvate accumulation was accompanied by a metabolic diversion leading to higher alanine abundance and reduced levels of TCA intermediates. Together, this comprehensive profiling provides a more detailed picture of the metabolic state. For instance, changes in less directly related metabolites were also seen, like the increase of histidine abundance. In this way, basic profiling of metabolic changes following *pdha-1* inhibition might uncover new (genetic) factors that contribute to metabolic adaptation in these circumstances. Such information is not only valuable for research in *C. elegans* per se, but could also be used to study processes where PDH is involved such as the aberrant preferential activation of glycolysis in cancer cells^31^ or the regulation of brown adipose tissue metabolism^32^. This is also the case for the metabolite changes observed upon *dpyd-1* RNAi in context to the chemotherapeutic drug 5-FU and for instance its toxicities in patients with DPYD variants^33^.

Similarly, when knocking down *elo-2* using RNAi, robust changes were observed in the total chain length of almost all lipid classes. Knockdown of *mdt-15* led to significant changes in the degree of saturation of a wide variety of lipid species. Combined, this shows that our method provides an exceptionally detailed view on complex lipid composition, which can be useful for the identification and study of disorders related to the lipidome.

RILs were used to illustrate that our method is capable of picking up not just large metabolic defects caused by knockdown of a single gene, but also more subtle metabolic effects in a non-interventional population harbouring genetic variation. While our study is not large enough to parse out the complexity of the underlying genetic traits, we clearly show both convergent and divergent patterns of inheritance, as expected based on population genetics. Future studies using the full panel of RILs will allow the reconstruction of genetic complexity that causes individual metabolic variation, and enable studying gene-by-environment interactions, i.e. which genes render the organism susceptible to environmental disturbances.

We also used the RILs to highlight examples of direct integration of lipidomic and metabolomic data. These multi-omics comparisons will be more meaningful when using our method, as both datasets originate from the same material, eliminating the noise of inherent biological differences when independent worm cultures are prepared. Integrating metabolomics and lipidomics data can be important, as polar and lipid pathways are interconnected and often converge in meaningful ways. As an example, we showed that the mitochondrial lipid CL correlates with the abundance of polar metabolites with important roles in mitochondrial function such as NAD^+^ and acetyl-CoA across the RIL panel. Such integration based on population data can support new hypotheses and its validation in natural populations.

On the technical side, the use of a diverse selection of internal standards allows for meaningful semi-quantitative comparisons between sample groups. Due to the ubiquitous and essential nature of many of the metabolites analyzed here, this dual extraction method can be developed to aid metabolomics and lipidomics in a wide variety of matrices. This is especially useful when the amount of available material is limited and two separate extractions might not be feasible, such as with human biopsy material or rare cell populations. When applying the current method to a new matrix, we strongly advise to perform a range-finding experiment to determine a sample quantity where most of the analytes of interest are in the linear range of the extraction and MS, as well as appropriate internal standards.

While the aforementioned considerations are important for all metabolite extraction methods, one of the main limitations of this method is that some polar lipid species are not ending up in the apolar layer but in the polar layer and are therefore not measured in lipidomics. When specifically interested in such polar lipid species, a dedicated one-phase lipidomics extraction yields a better result^20^. In conclusion, the currently presented method is capable of robustly analyzing a broad range of the metabolome and lipidome, and detecting biologically relevant differences while requiring only a single small sample.

## Methods

### Worm growth conditions for RNAi experiments and Recombinant Inbred Lines (RILs)

N2 worms and RIL strains were cultured at 20°C on nematode growth medium (NGM) agar plates seeded with OP50 strain *Escherichia coli*. For RNAi knockdown experiments, we seeded 2000 synchronized eggs per 10cm NGMi plate (containing 2□mM IPTG) with a bacterial lawn of either *E. coli* HT115 (RNAi control strain, containing an empty vector) or *pdha-1, dpyd-1, elo-2* or *mdt-15* RNAi bacteria. Similarly, for the parental strains (N2 and CB4856) and eight different offspring (RILs strains WN038, WN105, WN106, WN128, WN134, WN152, WN153, WN186), 2000 synchronized eggs were seeded per 10cm NGM plate, with a bacterial lawn of *E. coli* OP50. After 48 hours, the synchronous population at L4 larval stage was washed off the plates in M9 buffer and the worm pellet was washed with dH_2_O for three times and then collected in a 2□ml Eppendorf tube and snap frozen and stored at -80°C. Worm pellets were freeze-dried overnight and stored at room temperature until extraction.

### Two-phase extraction

In a 2 mL tube, the following amounts of internal standard dissolved in water were added to each sample of freeze dried worms for metabolomics: adenosine-^15^N_5_-monophosphate (5 nmol), adenosine-^15^N_5_-triphosphate (5 nmol), ^13^C_6_-fructose-1,6-diphosphate (1 nmol), guanosine-^15^N_5_-monophosphate (5 nmol), guanosine-^15^N_5_-triphosphate (5 nmol), ^13^C_6_-glucose (10 nmol), ^13^C_6_-glucose-6-phosphate (1 nmol), ^13^C_3_-pyruvate (0.5 nmol). In the same 2 mL tube, the following amounts of internal standards dissolved in 1:1 (v/v) methanol:chloroform were added for lipidomics: Bis(monoacylglycero)phosphate BMP(14:0)_2_ (0.2 nmol), Cardiolipin CL(14:0)_4_ (0.1 nmol), Lysophosphatidicacid LPA(14:0) (0.1 nmol), Lysophosphatidylcholine LPC(14:0) (0.5 nmol), Lysophosphatidylethanolamine LPE(14:0) (0.1 nmol), Lysophosphatidylglycerol LPG(14:0) (0.02 nmol), Phosphatidic acid PA(14:0)_2_ (0.5 nmol), Phosphatidylcholine PC(14:0)_2_ (0.2 nmol), Phosphatidylethanolamine PE(14:0)_2_ (0.5 nmol), Phosphatidylglycerol PG(14:0)_2_ (0.1 nmol), Phosphatidylserine PS(14:0)_2_ (5 nmol), Ceramide phosphocholine SM(d18:1/12:0) (2 nmol) (Avanti Polar Lipids, Alabaster, AL).

After adding IS mixes, a 5□mm steel bead and polar phase solvents (for a total of 500 µL water and 500 µL MeOH) were added and samples were homogenized using a TissueLyser II (Qiagen) for 5□min at a frequency of 30 times/sec. Chloroform was added for a total of 1 mL to each sample before thorough mixing. Samples were centrifuged for 10 minutes at 14,000 rpm. Of the two-phase system that was now created with protein precipitate in the middle, the top layer containing the polar phase was transferred to a new 1.5 mL Eppendorf tube. The bottom layer, containing the apolar fraction, was transferred to a 4 mL glass vial. The protein pellet in between the two layers was dried and subsequently dissolved in 0.2□M NaOH for quantification using a Pierce™ BCA Protein Assay following product protocol.

### One-phase lipidomic extraction

In a 2 mL tube, the following amounts of internal standards dissolved in 1:1 (v/v) methanol:chloroform were added to each sample: Bis(monoacylglycero)phosphate BMP(14:0)_2_ (0.2 nmol), Cardiolipin CL(14:0)_4_ (0.1 nmol), Lysophosphatidicacid LPA(14:0) (0.1 nmol), Lysophosphatidylcholine LPC(14:0) (0.5 nmol), Lysophosphatidylethanolamine LPE(14:0) (0.1 nmol), Lysophosphatidylglycerol LPG(14:0) (0.02 nmol), Phosphatidic acid PA(14:0)_2_ (0.5 nmol), Phosphatidylcholine PC(14:0)_2_ (0.2 nmol), Phosphatidylethanolamine PE(14:0)_2_ (0.5 nmol), Phosphatidylglycerol PG(14:0)_2_ (0.1 nmol), Phosphatidylserine PS(14:0)_2_ (5 nmol), Ceramide phosphocholine SM(d18:1/12:0) (2 nmol) (Avanti Polar Lipids, Alabaster, AL). After adding the IS mix, a steel bead and 1.5 mL 1:1 (v/v) methanol:chloroform were added to each sample. Samples were homogenized using a TissueLyser II (Qiagen) for 5□min at 30 Hz. Each sample was then centrifuged for 10 min at 14,000 rpm. Supernatant was transferred to a 4 mL glass vial.

### Metabolomics

After the polar phase was transferred to a new 1.5 mL tube, it was dried using a miVac vacuum concentrator at 60°C and processed as reported before^23^. The residue was dissolved in 100 µL 6:4 (v/v) methanol:water. Metabolites were analyzed using a Thermo Scientific Ultimate 3000 binary UPLC coupled to a Q Exactive Plus Orbitrap mass spectrometer. Nitrogen was used as the nebulizing gas. The spray voltage used was 2500□V, and the capillary temperature was 256□°C. S-lens RF level: 50, Auxilary gas: 11, Auxiliary gas temperature 300□°C, Sheath gas: 48, Sweep cone gas: 2. Samples were kept at 12°C during analysis and 5 µL of each sample was injected. Chromatographic separation was achieved using a Merck Millipore SeQuant ZIC-cHILIC column (PEEK 100 x 2.1 mm, 3 µm particle size). Column temperature was held at 30°C. Mobile phase consisted of (A) 1:9 (v/v) acetonitrile:water and (B) 9:1 (v/v) acetonitrile:water, both containing 5 mmol/L ammonium acetate. Using a flow rate of 0.25 mL/min, the LC gradient consisted of: 100% B for 0-2 min, reach 0% B at 28 min, 0% B for 28-30 min, reach 100% B at 31 min, 100% B for 31-32 min. Column re-equilibration is achieved by increasing the flow rate to 0.4 mL/min at 100% B for 32-35 min. MS data were acquired using negative ionization in full scan mode over the range of m/z 50-1200. Data were analyzed using Thermo Scientific Xcalibur software version 4.1.50. All reported metabolite intensities were normalized to appropriate internal standards, as well as total protein content in samples, determined using a Pierce™ BCA Protein Assay Kit. Metabolite identification has been based on a combination of accurate mass, (relative) retention times and fragmentation spectra, compared to the analysis of relevant standards.

### Lipidomics

After the solvents containing the lipids were transferred to a 4 mL glass vial, they were evaporated under a stream of nitrogen at 45°C. The residue was dissolved in 150 μL of 1:1 (v/v) chloroform:methanol. Lipids were analyzed using a Thermo Scientific Ultimate 3000 binary UPLC coupled to a Q Exactive Plus Orbitrap mass spectrometer. Nitrogen was used as the nebulizing gas. The spray voltage used was 2500□V, and the capillary temperature was 256□°C. S-lens RF level: 50, Auxilary gas: 11, Auxiliary gas temperature 300□°C, Sheath gas: 48, Sweep cone gas: 2. For normal phase separation, 2 μL of each sample was injected onto a Phenomenex® LUNA silica, 250 * 2 mm, 5µm 100Å. Column temperature was held at 25°C. Mobile phase consisted of (A) 85:15 (v/v) methanol:water containing 0.0125% formic acid and 3.35 mmol/L ammonia and (B) 97:3 (v/v) chloroform:methanol containing 0.0125% formic acid. Using a flow rate of 0.3 mL/min, the LC gradient consisted of: 10% A for 0-1 min, reach 20% A at 4 min, reach 85% A at 12 min, reach 100% A at 12.1 min, 100% A for 12.1-14 min, reach 10% A at 14.1 min, 10% A for 14.1-15 min. For reversed phase separation, 5 μL of each sample was injected onto a Waters HSS T3 column (150 x 2.1 mm, 1.8 μm particle size). Column temperature was held at 60°C. Mobile phase consisted of (A) 4:6 (v/v) methanol:water and B 1:9 (v/v) methanol:isopropanol, both containing 0.1% formic acid and 10 mmol/L ammonia. Using a flow rate of 0.4 mL/min, the LC gradient consisted of: 100% A at 0 min, reach 80% A at 1 min, reach 0% A at 16 min, 0% A for 16-20 min, reach 100% A at 20.1 min, 100% A for 20.1-21 min. MS data were acquired using negative and positive ionization using continuous scanning over the range of m/z 150 to m/z 2000. Data were analyzed using an in-house developed metabolomics pipeline written in the R programming language^34^. In brief, it comprises the following five steps: (1) pre-processing using the R package XCMS, (2) identification of metabolites, (3) isotope correction, (4) normalization and scaling and (5) statistical analysis^14^. All reported lipids were normalized to corresponding internal standards according to lipid class, as well as total protein content in samples, determined using a Pierce™ BCA Protein Assay Kit. Lipid identification has been based on a combination of accurate mass, (relative) retention times, and the injection of relevant standards.

### Heritability estimation of RILs and integration of metabolomics and lipidomics

Broad-sense heritability (*H*^2^) was calculated as described before^8^. Briefly, using an ANOVA explaining the metabolite variation over the offspring strains, the broad-sense heritability was calculated as *H*^2^ = V_g_ /(V_g_+V_e_), where *H*^2^ is the broad-sense heritability, V_g_ is the variance attributed to genetics and V_e_ is the variance attributed to other factors (e.g. measurement uncertainty or other biological factors). Significance of the heritability was calculated by permutation, where the trait values were randomly assigned to strains. Over these permutated values, the variance captured by strain and the residual variance were calculated. This procedure was repeated 1000 times for each trait. The obtained values were used as the by-chance distribution, and an FDR = 0.05 was taken as the 50th highest value. In the parental strains the broad-sense heritability was calculated as *H*^2^ = 0.5V_g_ /(0.5V_g_+V_e_), The factor 0.5 corrects for the overestimation of the additive variation in inbred strains^35^. The same permutation approach as for the broad-sense heritability was applied, taking the FDR = 0.05 threshold as significant. An *H*^2^ above the FDR value indates there is only a 5% chance the result is a false-positive.

Cross comparison between metabolites and lipids were performed as follows: Individual metabolite abundances were used. Lipid class abundances were calculated by summing the abundances of each lipid species from a given lipid class. These were cross-correlated using the imgCor function from the mixOmics package v6.6.2^36^ in R. Association and significance between lipids and metabolites was tested for using Pearson’s product moment correlation coefficient. Visualization of data was performed using ggplot2^37^. R v3.4.3 and Bioconductor v3.5 were used in these analyses.

## Acknowledgements

Work in the Houtkooper group is financially supported by an ERC Starting grant (no. 638290), a VIDI grant from ZonMw (no. 91715305), and a grant from the Velux Stiftung (no. 1063).

## Author contributions

MM, BVS, HLE, AWG, MATV, RLS, and MvW performed experiments. MLP, ACL, GEJ, and AHCvK performed bioinformatics. MM, BVS, RLS, MGS, JEK, GEJ, FMV, MvW and RHH interpreted data. MM, BVS, GEJ, MvW, and RHH wrote the manuscript with contributions from all others.

---

**Supplemental fig. S1.**
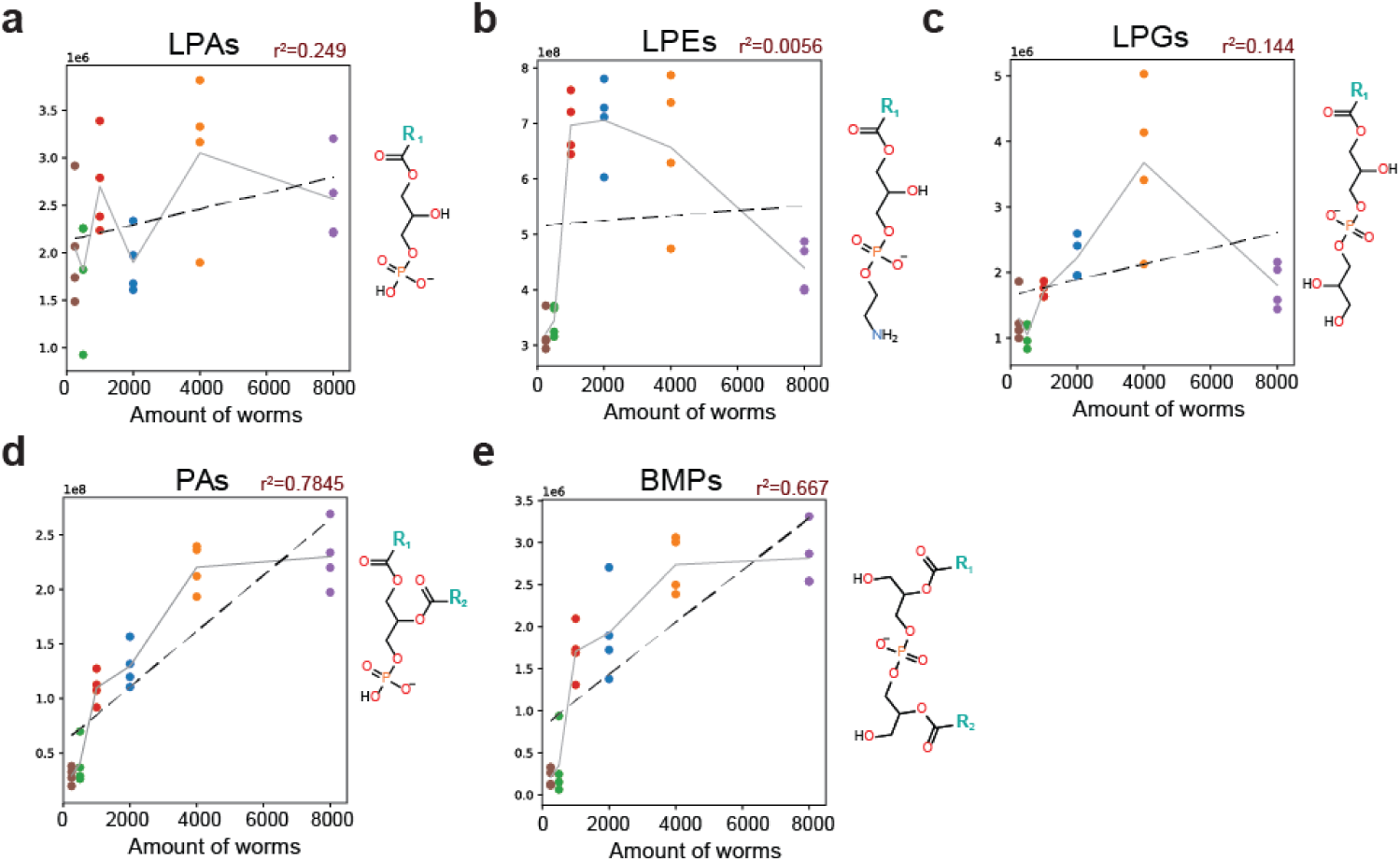
Validation of lipids extracted from apolar phase on the polar side of the lipid spectrum. Linearity of **a**, LPAa **b**, LPEs **c**, LPGs **d**, PAs **e**, BMPs. Chemical structure of the classes are depicted on the right of each graph.

**Supplemental fig. S2.**
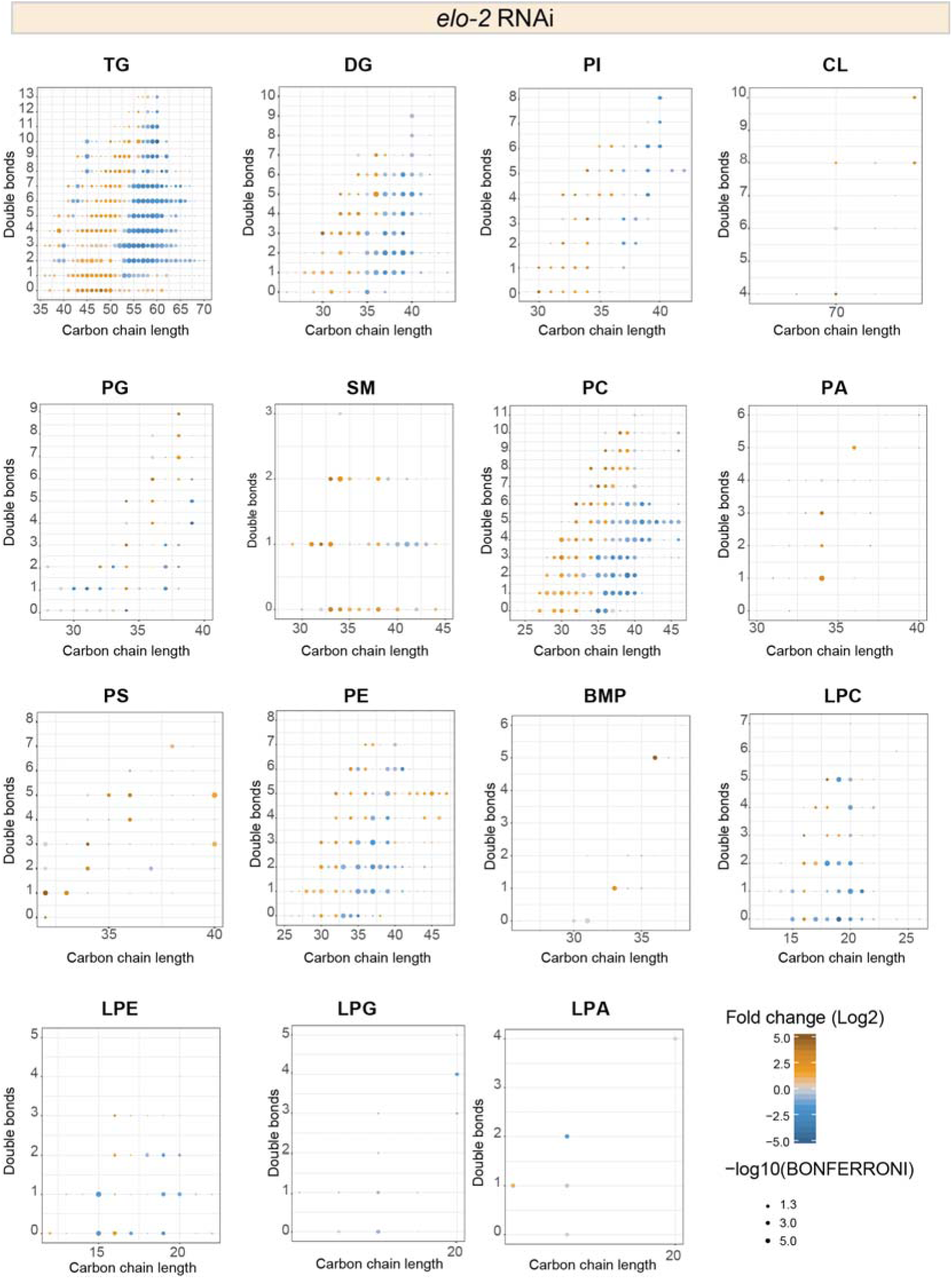
Phospholipid changes in all classes in worms treated with *elo-2* RNAi. Changes in the composition of almost all classes shows significant decrease of phospholipids with long carbon-chain length and significant increase of species with short carbon-chain length.

**Supplemental fig. S3.**
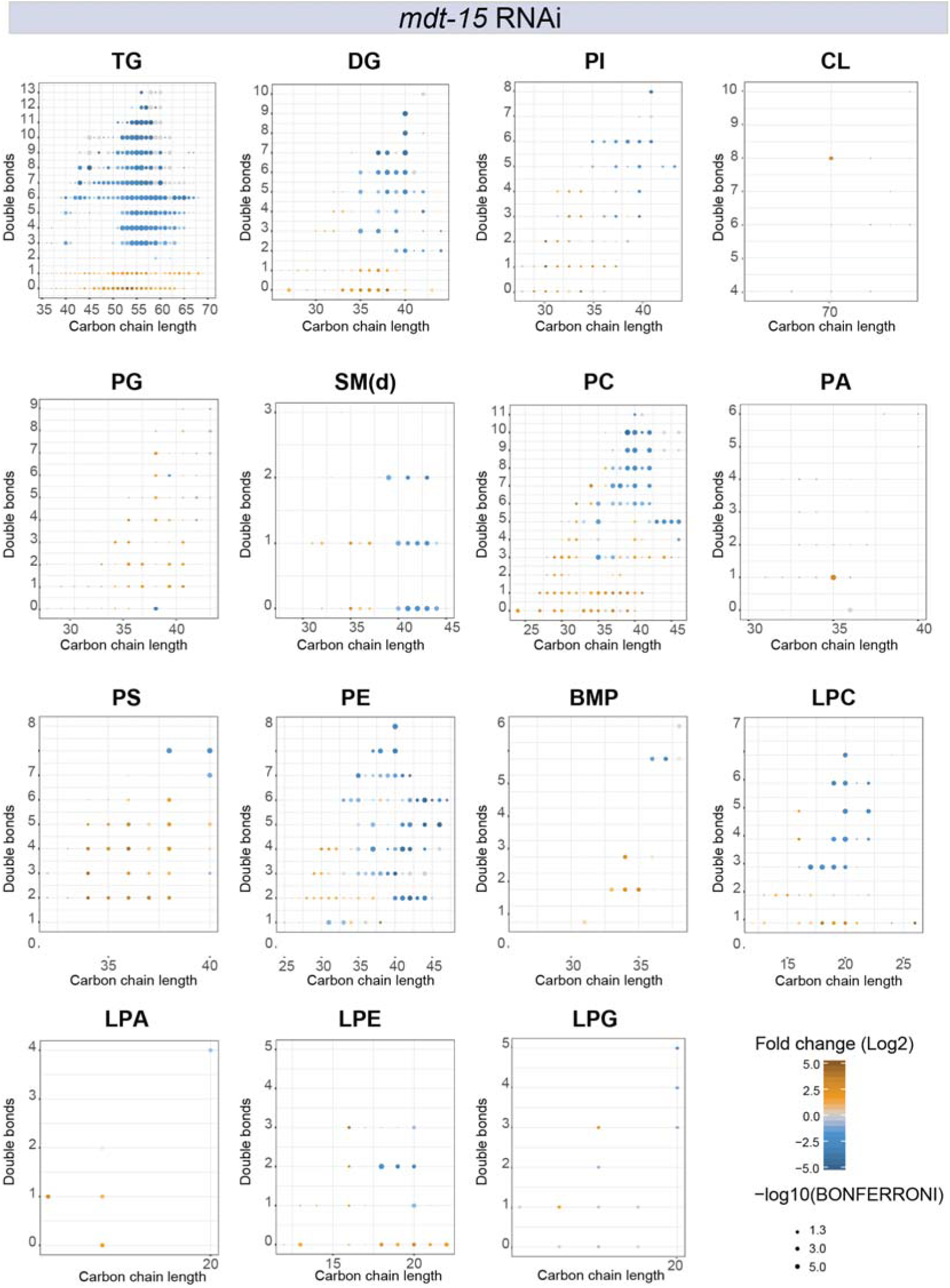
Phospholipid changes in all classes in worms treated with *mdt-15* RNAi. Changes in phospholipids of many species shows significant decrease of lipids with >2 double bonds and increase of PL with <2 double bonds.

